# Ankylosis homologue mediates cellular efflux of ATP, not pyrophosphate

**DOI:** 10.1101/2021.08.27.457978

**Authors:** Flora Szeri, Fatemeh Niaziorimi, Sylvia Donnelly, Nishat Fariha, Mariia Tertyshnaia, Drithi Patel, Stefan Lundkvist, Koen van de Wetering

**Affiliations:** Department of Dermatology and Cutaneous Biology, Jefferson Institute of Molecular Medicine and PXE International Center of Excellence in Research and Clinical Care, Sidney Kimmel Medical College, Thomas Jefferson University, Philadelphia (PA), USA; Research Centre for Natural Sciences, Institute of Enzymology, Budapest, Hungary

**Keywords:** ATP, anion transport, nucleoside/nucleotide transport, bone, plasma membrane, PPi, ecto-nucleotidases, ENPP1, ANKH/Ank, Ectopic mineralization.

## Abstract

The plasma membrane protein Ankylosis Homologue (ANKH, mouse ortholog: Ank) prevents pathological mineralization of joints by controlling extracellular levels of the mineralization inhibitor pyrophosphate (PPi). It was long thought that ANKH acts by transporting PPi into the joints, but we recently showed that ANKH releases large amounts of nucleoside triphosphates (NTPs), predominantly ATP, into the culture medium. This ATP is converted extracellularly into PPi and AMP by the ectoenzyme Ectonucleotide Pyrophosphatase Phosphodiesterase 1 (ENPP1). We could not rule out, however, that cells also release PPi directly via ANK. We now addressed this question by determining the effect of a complete absence of ENPP1 on ANKH-dependent extracellular PPi concentrations. Introduction of ANKH in ENPP1-deficient HEK293 cells resulted in robust cellular ATP release without the concomitant increase in extracellular PPi seen in ENPP1-proficient cells.

Ank-activity was previously shown to be responsible for about 75% of the PPi found in mouse bones. However, bones of *Enpp1*^*-/-*^ mice contained < 2.5% of the PPi found in bones of wild type mice, showing that Enpp1-activity is also a prerequisite for Ank-dependent PPi incorporation into the mineralized bone matrix *in vivo*. Hence, ATP release precedes ENPP1-mediated PPi formation. We find that ANKH also provides about 25% of plasma PPi, whereas we have previously shown that 60-70 % of plasma PPi is derived from the NTPs extruded by the ABC transporter, ABCC6. Both transporters that keep plasma PPi at sufficient levels to prevent pathological calcification, therefore do so by extruding NTPs rather than PPi itself.

## Introduction

The membrane protein ANKH (mouse ortholog: Ank) is a crucial inhibitor of joint calcification and its absence results in ankylosis in both humans and mice (1–3). Affected joints show massive ectopic calcification resulting in degradation of articular cartilage and ankylosis. For many years it was thought that ANKH prevents ectopic calcification by mediating cellular release of the calcification inhibitor inorganic pyrophosphate (PPi) into the joint space (2, 4, 5). Very recently we discovered that ANKH has an unexpected role in the extrusion of a range of cellular organic anions (6). These include malate, succinate and especially citrate, but also nucleoside triphosphates (NTPs). The released NTPs are extracellularly converted into PPi and their respective nucleoside monophosphate (NMP) by the ectonucleotidase pyrophosphatase/phosphodiesterase 1 (ENPP1). Our calculations indicated that at least 70% of the PPi present in the extracellular environment of ANKH-containing HEK293 cells came from cellular NTP release (6). We could not exclude the possibility, however, that a substantial part of the remaining 30% of the extracellular PPi was due to direct transport of PPi out of the cells by ANKH, as suggested in the older literature (2, 7). To evaluate whether cells release substantial amounts of PPi in an ANKH-dependent manner, we employed HEK293 cells deficient for ENPP1 (HEK293-*ENPP1*^*ko*^ cells), mice deficient in Ank (*Ank*^*ank/ank*^) and mice deficient in Enpp1 (*Enpp1*^*asj/asj*^). We find that release of NTPs, predominantly ATP, precedes formation of > 97% of the PPi found in the extracellular environment of ANKH-containing cells. This shows that ANKH does not transport substantial amounts of PPi out of cells, in contrast to the prevalent view within the field. Instead, ANKH mediates cellular release of ATP and other NTPs, which are converted into PPi by ENPP1 in the extracellular environment.

## Results and Discussion

### HEK293-ENPP1^ko^ cells degrade NTPs but are unable to generate extracellular NTPs

*ENPP1* was targeted in HEK293 cells using Crispr/Cas9. Clonal ENPP1 deficient cell lines were generated by sorting single cells into wells of 96-well plates. Two correctly targeted clones were selected for further analysis, HEK293-ENPP1^ko^ clone 3D5 and HEK293-ENPP1^ko^ clone 1D4 (Supporting Information Fig. S1). ENPP1 was readily detected by immunoblot analysis in HEK293-parental cells, but was undetectable in clones 3D5 and 1D4 (Fig. 1A). The ability to form PPi in the extracellular environment was determined in these lines by adding 20 μM ATP to the culture medium and quantifying extracellular PPi concentrations after 24 hours of incubation. Of note, serum-free Pro293a medium was used for these experiments, as serum is known to contain substantial amounts of a soluble form of ENPP1 (8). HEK293 parental cells formed about 9 μM PPi out of the 20 μM ATP added. Medium of both ENPP1^ko^ clones contained about 1 μM of PPi (Fig. 1B), levels that are similar to the background PPi concentrations detected in the Pro293 medium (indicated by the dashed line in Fig. 1B). These data show that the generated HEK293-ENPP1^ko^ cells almost completely lack the ability to form PPi from ATP added to the culture medium. Similar results were obtained when 20 μM GTP was added to the culture medium (Supporting Information Fig S2).

**Figure 1:**
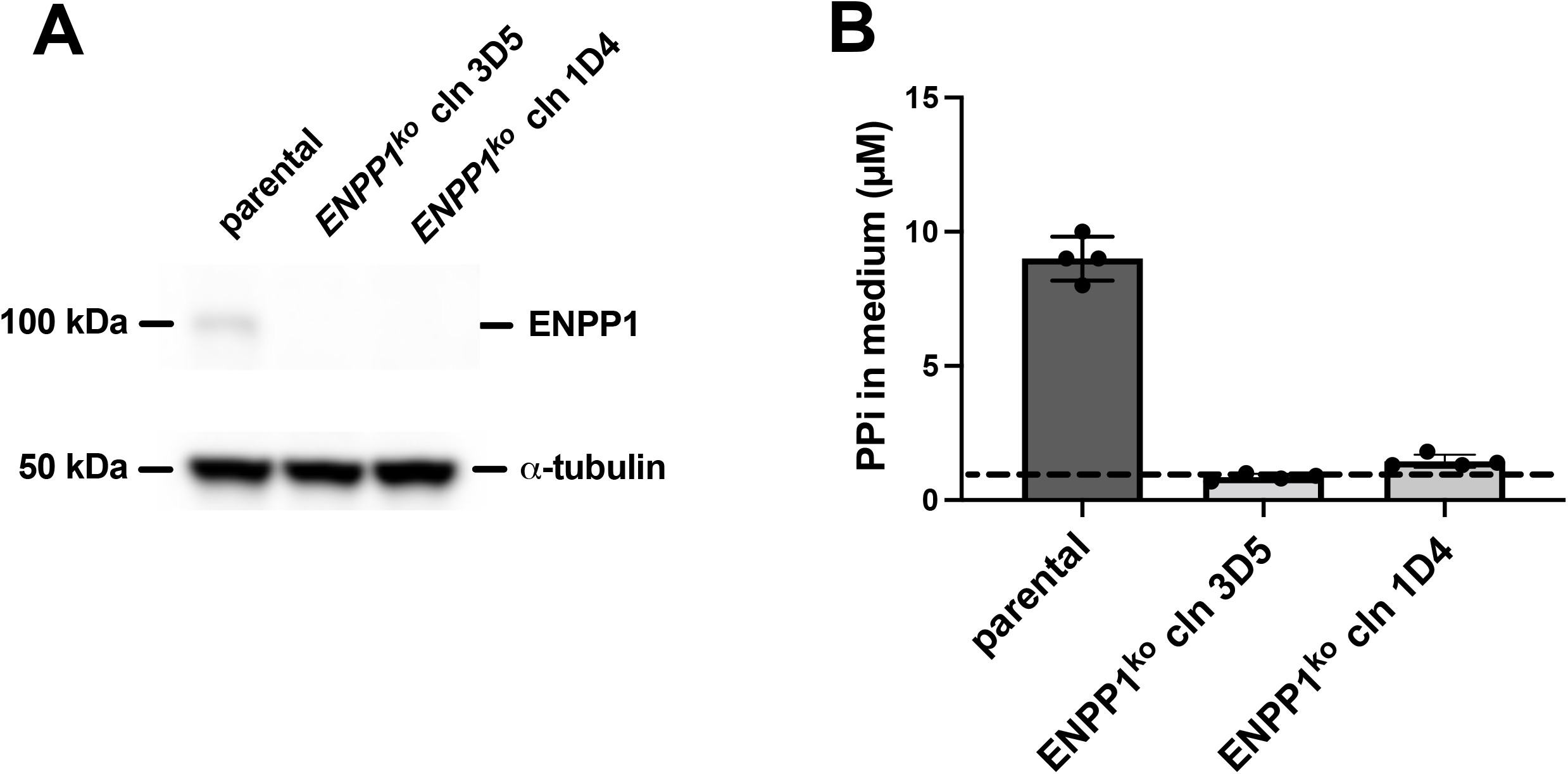
HEK293 cells lacking ENPP1 are severely hampered in their ability to convert extracellular ATP into pyrophosphate. *ENPP1* was targeted in HEK293 cells by Crispr/cas9 technology. In selected clonal cell lines, absence of ENPP1 was confirmed by immunoblot analysis (**A**). In (**B**), the indicated clonal cell lines were incubated for 24 hours in medium containing 20 μM ATP. The amount of pyrophosphate was subsequently quantified in collected 24-hour medium samples. A representative experiment repeated at least twice is shown in (**B**). Data are presented as mean ± SD of an experiment performed in quadruplicate (n=4). The dashed line in panel B indicates average concentrations of background PPi detected in culture medium not exposed to cells.

Our finding that a bit less than 50% of the ATP added to the medium samples of the ENPP1-proficient parental cells was converted into PPi, indicates that ENPP1 competes with other ecto-nucleotidases, like ectonucleoside triphosphate diphosphohydrolase (ENTPD1) (9), for available substrate. Twenty-four (24) hours after addition, no ATP was detected in medium of any of the 3 cell lines tested. These results imply efficient degradation, possibly by ENTPD1(9), of extracellular NTPs by HEK293 cells in the absence of ENPP1. Our finding that 4 hours after ATP addition medium of HEK293-ENPP1^ko^ cells contained similar concentrations of AMP as medium of the HEK293 parental cells (Supplemental Information, Fig. S3), provided further support for our conclusion that HEK293 cells efficiently degrade ATP, also in the absence of ENPP1.

### HEK293-ENPP1^ko^ cells overproducing ANKH^wt^ do not accumulate PPi in their culture medium

We selected HEK293-ENPP1^ko^ clone 3D5 for further analyses and generated HEK293-parental and HEK293-ENPP1^ko^ cells overproducing wild type ANKH (ANKH^wt^). As a control, HEK293-parental and HEK293-ENPP1^ko^ cells that overexpress the inactive ANKH mutant, ANKH^L244S^ (1), were used. Figure 2A displays ANKH immunoblot analysis, showing roughly similar protein levels for ANKH^wt^ and ANKH^L244S^ in the cell lines generated (Fig. 2A). A real-time ATP efflux assay showed robust ATP efflux from HEK293 cells overproducing ANKH^wt^ (Fig. 2B), independent of the presence of ENPP1 (Fig. 2B). Absence of ENPP1 in HEK293-ANKH^wt^ cells did not result in higher ATP levels in the real-time ATP efflux assay. This indicates that released ATP is also rapidly degraded in the extracellular environment in the absence of ENPP1. As expected (6), cells containing the inactive L244S ANKH mutant (1, 6) did not release substantial amounts of ATP into the extracellular environment (Fig. 2B).

**Figure 2:**
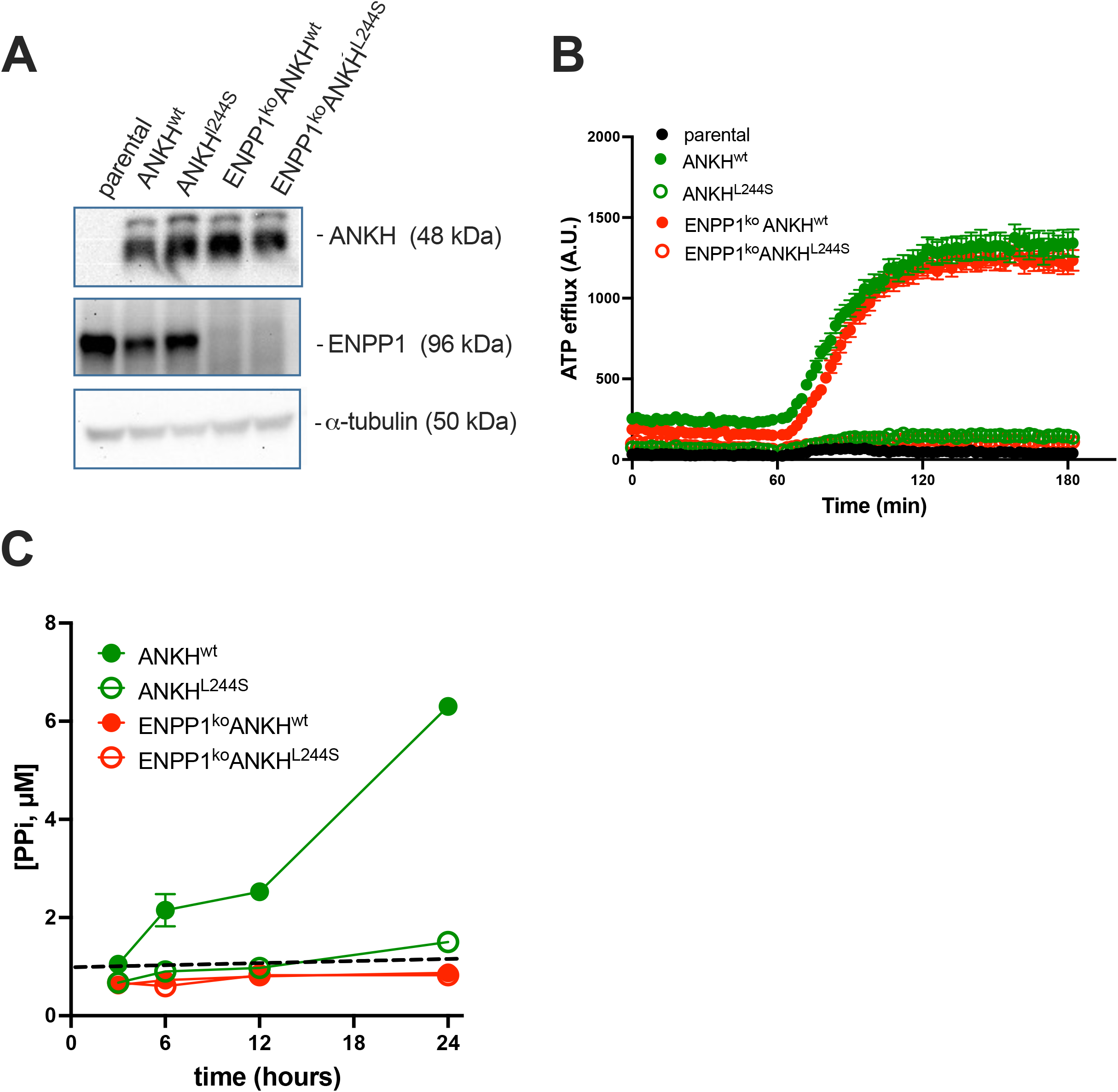
HEK293 cells need both ENPP1 and ANKH to accumulate PPi in their culture medium. (**A**) Immunoblot analysis demonstrates similar amounts of ANKH protein in HEK293 cells overproducing ANKH^wt^ and the inactive ANKH^L244S^ mutant. Absence of ENPP1 was confirmed in the ENPP1^KO^ cell lines. α-Tubulin was used as a loading control. (**B**) ATP efflux from the indicated cell lines grown till confluent in 96-well plates was followed in real time using luciferase/luciferin. Data of a representative experiment repeated at least 4 times is shown and represent mean ± SEM (n=16). (**C**) The indicated HEK293 clones were incubated for 24 hours in serum-free Pro293A medium. Samples were taken from the culture medium for pyrophosphate analysis at the indicated time points and analyzed for pyrophosphate (PPi) concentration. Data of a representative experiment repeated at least twice is shown and represent mean ± SD of an experiment performed in quadruplicate. The dashed line in panel C indicates average concentrations of PPi detected in culture medium not exposed to cells.

Next, we followed PPi accumulation in the culture medium over time (Fig. 2C). In the absence of ENPP1, PPi levels hardly increased during the 24-hour incubation period, independent of the presence of ANKH^wt^. In ENPP1-proficient cells, however, we detected clear accumulation of PPi in the culture medium of ANKH^wt^-containing cells over time. These data show that the PPi that accumulates in medium of ANKH^wt^-containing HEK293 cells depends on the presence of ENPP1 and that, at least in HEK293 cells, direct PPi efflux does not substantially contribute to the ANKH-dependent accumulation of PPi in the extracellular environment.

### NTP release underlies Ank-dependent deposition of PPi in bone

PPi is predominantly known for its role in prevention of soft tissue mineralization (10, 11). Intriguingly, the mineral phase of bone contains large amounts of PPi (6, 12). Ank is highly expressed in osteoblasts(13) and a major determinant of PPi incorporation in bone (6). To explore if NTP release also underlies Ank-dependent extracellular PPi deposition *in vivo*, we determined the amount of PPi present in femora and tibiae of *Enpp1*^*-/-*^ mice (Fig. 3). Bones of *Enpp1*^*-*^ deficient mice contained less than 2% of the PPi found in bones of control animals. We previously found that 75% of the PPi in tibiae and femora depend on Ank activity (Fig. 3A, *Ank*^*ank/ank*^ data taken from (6)). The almost complete absence of PPi in tibiae and femora of *Enpp1*^*-/-*^ mice implies that Enpp1-mediated conversion of extracellular ATP is required for close to all of the PPi in the mineral phase of bone. This means that also the 75% of PPi in bone that depends on Ank activity (Fig. 3A) must originate from Enpp1-dependent conversion of extracellular ATP. Also in bone *in vivo*, ATP release clearly precedes Ank-mediated PPi accumulation, providing additional evidence that cells do not release substantial amounts of PPi via Ank. Direct Ank-dependent PPi release can be calculated to be responsible for at most 2.5% of the PPi ending up in bone matrix: the 2% detected in bones of *Enpp1*^*-/-*^ mice divided by the 75% of PPi found in bone that depends on Ank-activity. This is an overestimation of the potential contribution of direct transport of PPi by Ank, however, as the *Enpp1*^*-/-*^ mouse strain used for our studies (Enpp1^asj^) is a hypomorph, which still shows some Enpp1 activity (14). In addition, the removal of bone marrow from the bone tissue is never complete. Some cells will remain in the bone matrix and their intracellular PPi might somewhat contribute to the amounts of PPi detected in the bone extracts. In conclusion, analysis of bones of *Enpp1*^*-/-*^ and *Ank*^*ank/ank*^ mice, recapitulates our results obtained in HEK293 cells, and shows that also in intact animals, Ank does not mediate release of substantial amounts of PPi from cells.

**Figure 3:**
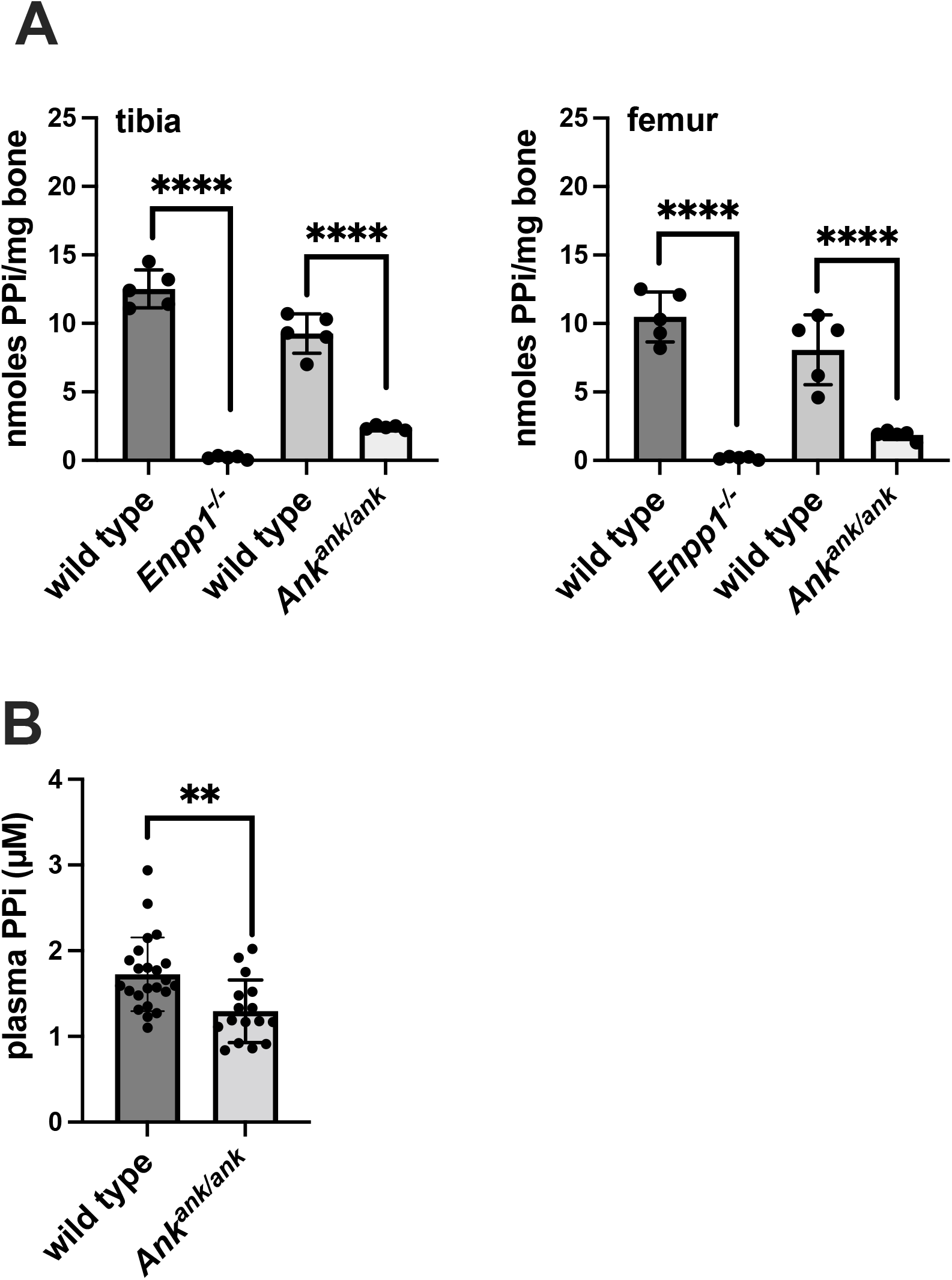
Pyrophosphate (PPi) amounts in bones of *Enpp1*^*-/-*^ and *Ank*^*ank/ank*^ mice and in plasma of *Ank*^*ank/ank*^ mice are reduced, confirming absence of substantial cellular PPi release via Ank *in vivo*. (**A**) Pyrophosphate is virtually absent in tibiae and femora of *Enpp1*^*-/-*^ mice and reduced by over 75% in bones of *Ank*^*ank/ank*^ mice. *Enpp1*^*-/-*^ (C57Bl6/J) and *Ank*^*ank/ank*^ (C3FeB6) mice were on a different genetic background, which might explain the slightly different amounts of PPi detected in bones of both wild type strains. Differences did not reach statistical significance, however. Data represent mean ± SD (n=5). (**B**) Plasma concentrations of PPi in wild type (n=23) and *Ank*^*ank/ank*^ (n=16) mice. Similar numbers of male and female mice were included in these experiments. Data are presented as mean ± SD.

### Plasma of *Ank*^*ank/ank*^ mice contains reduced concentrations of PPi, compatible with a role of Ank in NTP release

It has previously been demonstrated that plasma of *Enpp1*^*-/-*^ mice is virtually devoid of PPi (15). This implies that NTP release precedes all the PPi present in plasma. To provide additional evidence that Ank is involved in NTP release and not of PPi, we determined plasma PPi concentrations in wild-type and *Ank*^*ank/ank*^ mice. About 60-70% of the PPi present in plasma depends on ABCC6-dependent NTP release. Ank can therefore be expected to be responsible for maximally 30-40% of the PPi present in plasma. Our analyses indicated PPi concentrations in plasma of *Ank*^*ank/ank*^ mice were about 25% lower than levels detected in their wild type littermates (wild type: 1.7 ± 0.4 μM vs *Ank*^*ank/ank*^ 1.3 ± 0.4 μM, Fig. 3B), in line with recent data of Fujii et al. (4). As all PPi detected in plasma depends on ENPP1 (16), the fraction (~25%) that depends on Ank-activity, must be due to NTPs released into the circulation and cannot be explained by direct, Ank-dependent cellular efflux of PPi. This provides additional evidence that ANKH is involved in release of NTPs, not of PPi. It is important to note that Abcc6 and Ank both are involved in the cellular release of NTPs (6, 17, 18) and our data now indicate that together both membrane proteins account for the bulk of PPi present in plasma. Both proteins inhibit pathological mineralization by regulating extracellular PPi concentrations, though in different tissues: Absence of functional ABCC6 is associated with pseudoxanthoma elasticum (PXE), a rare hereditary mineralization disorder (10). Absence of ABCC6-mediated hepatic ATP release in PXE patients results in reduced PPi plasma concentrations and, as a consequence, progressive mineralization of blood vessels, skin and eyes. Currently there is no effective treatment for PXE and ectopic calcification therefore slowly progresses after diagnosis (19). Absence of ANKH/Ank also results in pathological mineralization. In ANKH-deficient individuals joint spaces of hand and feet and spine are mineralize, resulting in a severe form of ankylosis (1). Our finding that Ank also contributes to plasma PPi concentrations, makes this membrane protein an attractive target in PXE. Possibly, ANKH can be pharmacologically stimulated in PXE patients, to increase NTP release into the blood circulation for subsequent *in situ* PPi formation and normalization of plasma PPi concentrations. Increased ANKH-dependent release of the calcium chelator citrate, which also leaves cells in an ANK-dependent manner (6), might further contribute to inhibition of ectopic mineralization.

Because of its involvement in raising extracellular PPi concentrations, earlier reports described ANKH as a PPi efflux transporter(1, 2, 5, 7). To the best of our knowledge, no direct biochemical evidence that ANKH mediates cellular PPi efflux has been published up until know, however. Gurley and coworkers in Xenopus laevis oocytes overproducing ANKH found PPi, uptake, not efflux of PPi (7), suggesting that ANKH allows for bidirectional transport. Low intracellular PPi concentrations might have resulted in uptake of some extracellular PPi into the ANKH producing oocytes. This explanation is compatible with our current results as it implies that under normal conditions intracellular citrate, ATP and other NTPs by far outcompete PPi for release via ANKH. Last but not least, our finding that both ANKH and ENPP1 are needed to provide the extracellular environment with PPi also provides an explanation why Gurley et al. did not find increased levels of PPi in the extracellular milieu of their ANKH-producing oocytes (7): Xenopus laevis oocytes might just not express ENPP1 and hence are unable to convert released ATP into PPi.

In conclusion, we show that the PPi found in the extracellular environment of ANKH-containing cells is explained by cellular release of NTPs release, which are subsequently converted into NMPs and PPi by the ecto-nucleotidase ENPP1. Under physiological conditions ANKH does not mediate significant cellular PPi release, in contrast to the view commonly held in the calcification field.

## Experimental procedures

### Reagents

Unless otherwise indicated all reagents were obtained from Fisher Scientific (Waltham, MA).

### Cell culture

HEK293 cells were passaged in HyClone DMEM (GE Healthcare Systems, Marlborough, MA) supplemented with 5% FBS and 100 units pen/strep per ml (Gibco, Thermo Fisher, Waltham, MA) at 37°C and 5% CO_2_ under humidified conditions. Efflux experiments were performed in 6-well plates. 500,000 cells were seeded per well and 2 days later the experiment was started by replacing the culture medium with 2.5 ml Pro293a medium (Lonza, Basel, Switzerland), supplemented with 2 mM L-glutamine and 100 units pen/strep (Gibco, Thermo Fisher, Waltham, MA) per ml. Samples were taken at the indicated time points. At the end of the experiment, the presence of similar numbers of cells was confirmed by sulphorhodamine (SRB) staining (20).

### Generation of HEK293-ENPP1^ko^ cells

A pool of HEK293 cells deficient in ENPP1 was obtained from Synthego Corporation (Redwood City, CA) using proprietary technology (https://www.synthego.com/resources/all/protocols). In short, a protein mixture containing the Cas9 protein and the synthetic chemically modified single-guide RNA (GAUGGAGCGCGACGGCUGCG) were electroporated into the cells. Single cells were sorted in into 96-well plates using an BD FACSMelody cell sorter (BD Biosciences, Franklin Lakes, NJ). Correctly targeted clones were identified by Sanger sequencing.

### Generation of HEK293 control and HEK293-ENPP1^ko^ overexpressing ANKH^wt^ or the inactive ANKH mutant, ANKH^L244S^

HEK293 control and HEK293-ENPP1^ko^ cells were transfected with the pQCXIP expression vector, containing cDNAs encoding wild type ANKH (ANKH^wt^) or an inactive ANKH mutant, (ANKH^L244S^) as previously described (6). ANKH expression was determined in clones resistant to 2 μM puromycin (Gibco, Thermo Fisher, Waltham, MA) by immunoblot analysis with a polyclonal antibody directed against ANKH (OAAB06341, Aviva Systems Biology, San Diego, CA) and using α-tubulin as a loading control (mouse anti-α-tubulin B-5-1-2, Santa Cruz Biotechnology, Dallas, TX).

### Determination of ENPP1 activity

500,000 cells/well were seeded in 6-well plates. Two days later, the 5% FCS-containing DMEM culture medium was replaced by 2.5 ml serum-free Pro293a medium, with or without 20 μM ATP (Thermo Fisher, Waltham, MA). Of note, serum-free medium was used in these experiments as serum is known to contain a soluble form of ENPP1 (8). At the indicated time points, 100 μl samples were taken for PPi analysis. At the end of the experiment, relative cell density was determined using SRB staining (20).

### Real-time ATP efflux assays and detection of AMP in cell culture medium

ATP efflux from HEK293 cells was followed in real time as described (6, 17, 20). AMP was detected in culture medium as described in (17), with modifications. Samples were diluted 25-fold in Tris-EDTA (100 mM Tris pH 7.75, 2 mM EDTA) prior to analysis. To 500 μl of assay mixture (20 nM PPi, 20 nM phosphoenolpyruvate (PEP) and 29 mU pyruvate phosphate dikinase (PPDK, Kikkoman, Biochemifa Company, Noda City, Japan) in SRB buffer (BioThema, Handen, Sweden)), 5 μl of diluted medium sample was added. In the assay mixture, AMP, PPi and PEP were converted into ATP by PPDK, which resulted in an increase in luminescence generated by the SL-reagent. A known amount of ATP was finally added as internal standard and the ratio between the increase in bioluminescent signal induced by the addition of AMP and the internal ATP standard was used to calculate the PPi concentration of the sample. The assay was calibrated with 1 μM AMP and performed in a Berthold FB12 luminometer (Berthold Technologies, Bad Wild Bad, Germany) in the linear range of the detector.

### Animals

The *Enpp1*^*-/-*^ (*Enpp1*^*asj*^, official name: C57BL/6J-Enpp1asj/GrsrJ) mice were originally obtained from The Jackson Laboratory (stock No: 012810). Heterozygote breeders were used to generate *Enpp1*^*-/-*^ and wild type mice. Bones collected from leftover carcasses of female *Enpp1*^*-/-*^ animals (age range: 3-5 months) used in other experiments were used for bone PPi analyses. Mice heterozygous for the progressive ankylosis allele (*ank*) were obtained from The Jackson Laboratory (Bar Harbor, ME; C3FeB6 *A/A*^*w-J*^-*Ank*^*ank/J*^, stock number 000200). Heterozygote breeders were used to generate *Ank*^*ank/ank*^ and wild-type animals. Animals analyzed were between 2-3 months old at the time of blood sampling. Blood was collected by cardiac puncture in syringes containing 500 μl of ice-cold stop solution (21, 22), which consisted of 118 mM NaCl, 5 mM KCl, 40 mM tricine pH 7.4, 4.15 mM EDTA, 10 μM forskolin (to stabilize platelets), 100 μM isobutylmethylxanthine (IBMX, to stabilize platelets) and 5 nM S-4-nitrobenzyl-6-thioinosine (NBTI, to prevent ATP release by erythrocytes). Approximately 500 μl of blood was collected and the blood/stop solution mixture was emptied in pre-weighted 1.5 ml tubes to quantify the amount of blood collected. Plasma was prepared by centrifugation (2 min, 4 °C, 13000xg). Six hundred (600) μl plasma was transferred to a new tube and spun again (2 min, 4 °C, 13000xg) to remove any residual erythrocytes. Five hundred (500) μl plasma was subsequently stored at −80 °C until analysis. Studies on plasma PPi concentrations included similar numbers of male and female mice.

Animal studies were approved by the Institutional Animal Care and Use Committee of Thomas Jefferson University in accordance with the National Institutes of Health Guide for Care and Use of Laboratory Animals under approval number 02081 (*Ank*^*ank/ank*^ mice) and 00123 (*Enpp1*^*-/-*^ mice). To limit the number of animals used for our experiments, bones of the *Enpp1*^*-/-*^ mice were obtained from leftover carcasses from animals used in other experiments in which only female mice were used.

### Quantification of PPi in plasma, bone and medium samples

To quantify PPi in bone we used ATP sulfurylase to convert PPi into ATP in the presence of excess adenosine 5’ phosphosulfate (APS) and subsequent detection of ATP by luciferase/luciferin. PPi was determined in the plasma/stop solution mixture as previously described (17, 18), with modifications: to 10 μl of the plasma/stop solution 70 μl of mixture consisting of 75 mU/ml ATP sulfurylase (New England Biolabs, Cambridge, MA), 1 μmol/L APS (Santa Cruz Biotechnology, Dallas, TX), 80 μmol/L MgCl_2_ and 50 mmol/L HEPES (pH 7.4). After incubation for 30 min at 30 °C and 10 min at 90 °C, 10 μl of the reaction mixture was incubated with 30 μl of BactiterGlo (Promega, Madison, WI). Luminescence was determined using a Flex Station 3 plate reader (Molecular Devices, San Jose, Ca). Plasma concentrations were finally calculated by taking along a PPi standard curve and assuming a hematocrit of 40%.

PPi amounts in bone were determined as previously described (6). In short, 1000 μl 10% formic acid was added per 25 mg of bone from which bone marrow was removed by centrifugation. Samples we subsequently incubated overnight at 60 °C. After centrifugation (30,000 RCF, 4 °C, 10 min), the supernatant was stored at −80 °C until analysis. Bone extracts were diluted 500-fold in Tris-EDTA (100 mM Tris pH 7.75, 2 mM EDTA) buffer prior to analysis. PPi was quantified using 500 μl of the assay mixture to which 5 μl of (diluted) sample was added. In the assay mixture, PPi and APS were converted into ATP by ATPS, which resulted in an increase in luminescence. A known amount of ATP was finally added as internal standard and the ratio between the increase in bioluminescent signal induced by the addition of PPi and the internal ATP standard was used to calculate the PPi concentration of the sample. The assay was performed in a Berthold FB12 luminometer (Berthold Technologies, Bad Wild Bad, Germany) in the linear range of the detector.

## Data availability

All data are contained within the manuscript or in the supporting information.

## Supporting information

This article contains supporting information.

## Acknowledgements

We thank our colleagues, Qiaoli Li and Douglas Ralph (Both Thomas Jefferson University) for providing bones of *Enpp1*^*-/-*^ mice and Jouni Uitto (Thomas Jefferson University) and Piet Borst (The Netherlands Cancer Institute) for their critical evaluation of our manuscript.

## Funding and additional information

This research was funded by National Institutes of Health, Grant R01AR072695 (K.v.d.W.), U.S. Department of State (Fulbright Visiting Scholar Program), National Research, Development and Innovation Office (OTKA FK131946), Hungarian Academy of Sciences (Bolyai János Fellowship BO/00730/19/8, Mobility grant) and the Ministry for Innovation and Technology from the source of the National Research, Development and Innovation Fund (ÚNKP-2020 New National Excellence Program) to F.S. Further funding for this work was provided by PXE International for K.v.d.W. and F.S. Generation of ENPP1^ko^ HEK293 cell lines was financially supported by an intramural grant of Thomas Jefferson University. The content is solely the responsibility of the authors and does not necessarily represent the official views of the National Institutes of Health.

