## Supplemental Figures 1-3 for "Ankylosis homologue mediates cellular efflux of ATP, not pyrophosphate"

### Figure S1

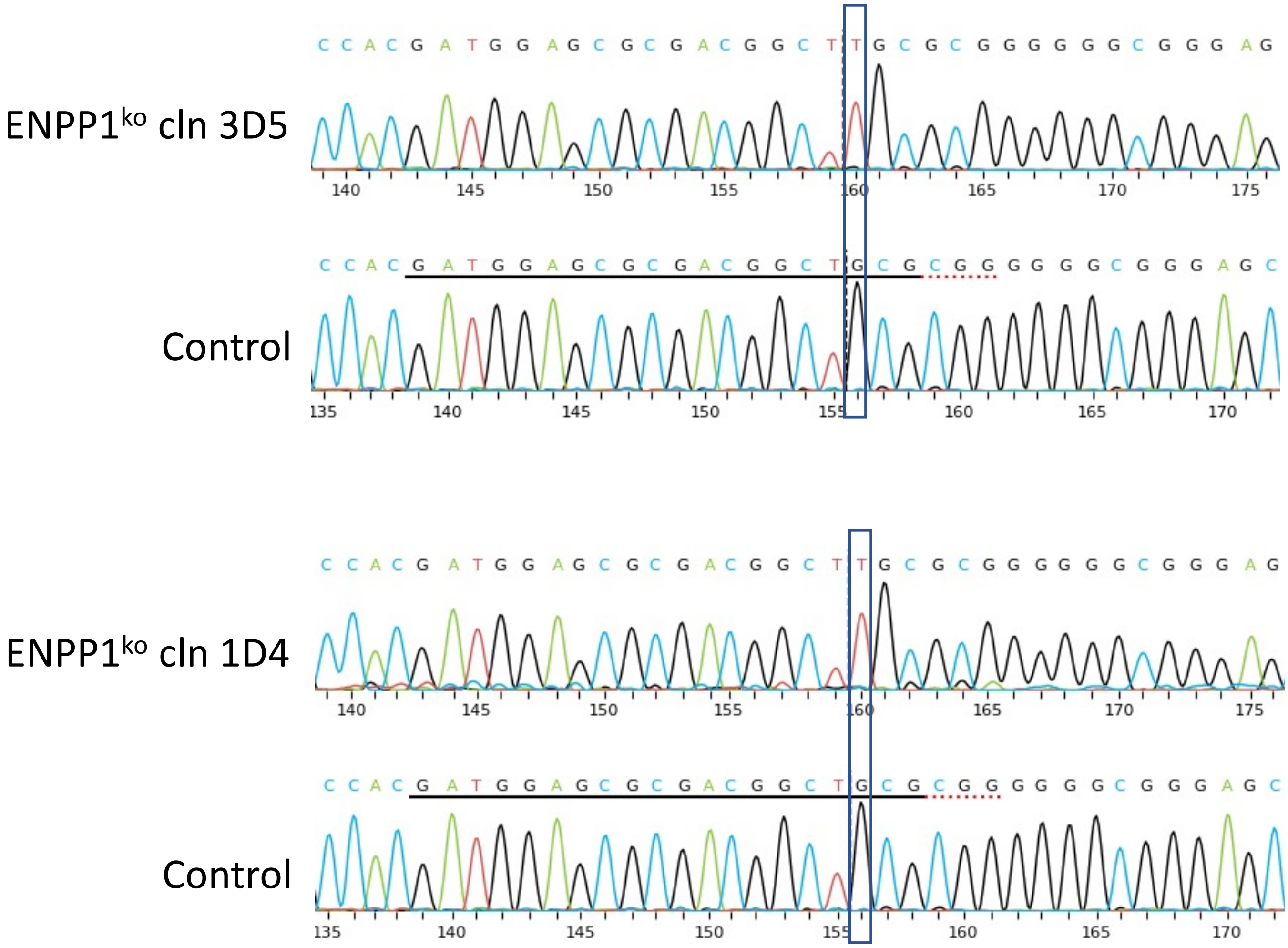


**Figure S1. Sanger sequencing shows homozygous insertion of a thymidine (T) in exon 1 of the *ENPP1* gene after Crispr/Cas9 targeting**. The guide sequence used for Crispr/Cas9 targeting is indicated by the bold black line in the control sequence. The online ICE v2 CRISPR analysis tool of Syntego (Redwood City, Ca) was used to analyze sequencing results and indicated 99 and 98% likeliness of a complete knockout for ENPP1ko cln 3D5 and ENPP1 cln 1D5, respectively.

### Figure S2

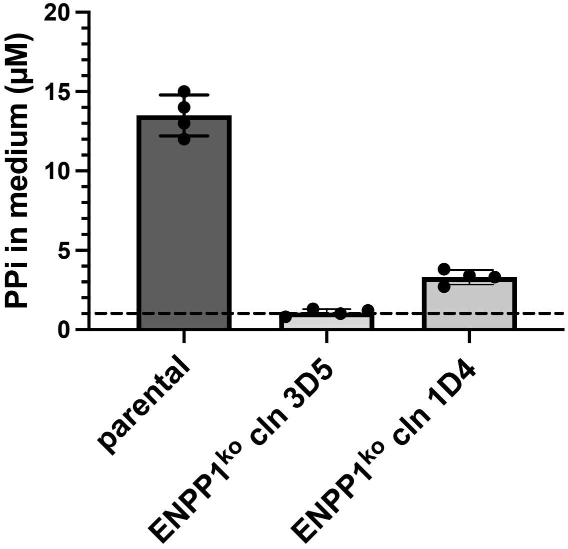


**Supplementary Information Figure S2. HEK293 cells lacking ENPP1 are severely hampered in their ability to convert extracellular GTP into pyrophosphate.** HEK293 cell lines proficient (parental) or deficient (ENPP1$^{ko}$ cln 3d5 and ENPP1$^{ko}$ cln 1D4) for ENPP1 were incubated in medium containing 20 $\mu$M GTP. Twenty-four (24) hours later, medium was collected and analyzed for pyrophosphate concentration as described in "Experimental Procedures". Data represent mean ± SD of an experiment performed in quadruplicate. The dashed line indicates average concentrations of background PPi detected in culture medium not exposed to cells.

### Figure S3

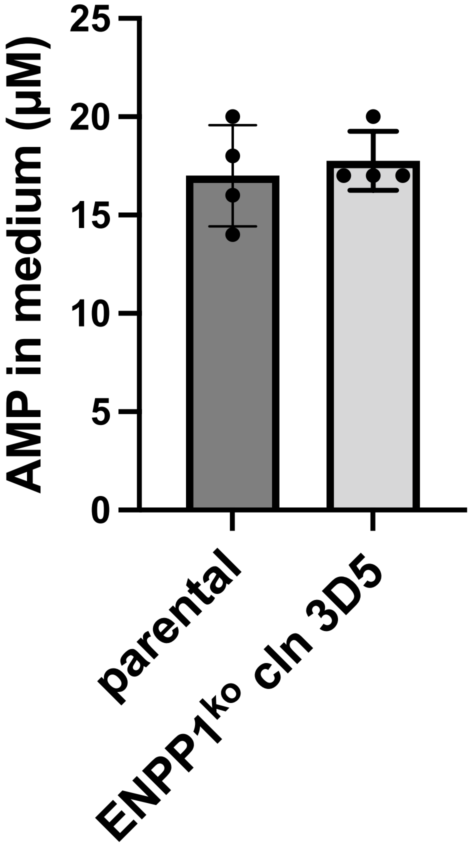


**Supporting Information Figure S3. Cells deficient in ENPP1 efficiently generate AMP from exogenously added ATP.** HEK293 cell lines proficient (parental) or deficient (ENPP1$^{ko}$ cln 3D5) for ENPP1 were incubated in medium containing 20 $\mu$M ATP. Four (4) hours later, AMP was quantified in the culture medium as described in “Experimental Procedures”. Data represent mean ± SD of an experiment performed in quadruplicate.
